# Metabolic and Anti-Proliferative Responses of Pancreatic Cancer Cells to Ultrasound and Nanobubble Treatment

**DOI:** 10.64898/2026.04.24.720507

**Authors:** Sila Appak-Baskoy, Muhammad Saad Khan, Farkhondeh Ghaderi, Agata A. Exner, Michael C. Kolios, Imogen R. Coe

**Affiliations:** Department of Chemistry and Biology, Toronto Metropolitan University, Toronto, ON M5B 2K3, Canada; Institute for Biomedical Engineering, Science and Technology (iBEST), Li Ka Shing Knowledge Institute, St. Michael’s Hospital, Toronto, ON M5B 1T8, Canada; Department of Physics, Toronto Metropolitan University, Toronto, ON M5B 2K3, Canada; Keenan Research Center for Biomedical Science, St. Michael’s Hospital, Unity Health Toronto, Toronto, ON M5B 1T8, Canada; Departments of Radiology and Biomedical Engineering, Case Western Reserve University School of Medicine, Cleveland, Ohio 44106, United States

**Keywords:** Nanobubble, Ultrasound, Theranostics, Proliferation, Metabolism

## Abstract

Pancreatic ductal adenocarcinoma (PDAC) remains one of the most lethal malignancies due to its dense stroma, which limits drug delivery and therapeutic efficacy. Ultrasound (US) mediated strategies using nanobubbles (NBs) offer a promising approach to enhance treatment, yet the biological effects of NB exposure and the timing of US application remain unclear. Here, we investigated how NB exposure with immediate (0h) or delayed (1h) US affects viability, proliferation, metabolism, and stress signaling in PANC-1 and BxPC-3 cells. Immediate US exposure in the presence of extracellular nanobubbles resulted in a greater reduction in cell viability at 24 h compared to delayed US application. Proliferation analysis showed that Ki67 positivity decreased following USNB treatments in both cell lines. Metabolically, NB treatment alone increased cellular activity, whereas combined USNB treatment reduced metabolic activity over time. Seahorse analysis revealed higher basal respiration in PANC-1 cells compared to BxPC-3 cells, consistent with a more glycolytic phenotype, while USNB treatment enhanced glycolytic responses, particularly in PANC-1. Moreover, stress responses were also more pronounced in PANC-1 cells, with HSP70 expression increasing up to 2-fold in NB incubated group and decreasing in USNB groups compared to untreated, whereas BxPC-3 cells exhibited only modest and opposite changes to PANC-1 in HSP70 expression decreasing with NB incubation. Treatment timing critically influenced outcomes, with immediate US producing stronger antiproliferative and cytotoxic effects, highlighting the importance of sequencing in USNB therapeutic strategies. Moreover, NBs alone stimulated metabolic and stress responses that may promote proliferation, whereas NBs combined with US induced stronger stress responses associated with metabolic reprogramming and reduced proliferation.

## Introduction

Pancreatic ductal adenocarcinoma (PDAC) remains one of the most lethal malignancies worldwide, with a five-year survival rate below 10% and minimal improvement in patient outcomes over the past several decades. The poor prognosis is largely attributed to late diagnosis, aggressive tumor biology, and limited therapeutic efficacy of current treatments. Standard therapeutic approaches, including surgical resection, chemotherapy, and radiotherapy, are often ineffective due to the unique tumor microenvironment of PDAC with the presence of an extensive desmoplastic stroma composed of cancer-associated fibroblasts, extracellular matrix proteins, and dense fibrotic tissue surrounding tumor cells which severely limits drug delivery and therapeutic penetration into tumor tissue (Sarantis et al, 2020). This stromal barrier increases interstitial fluid pressure, compresses tumor vasculature, reduces perfusion, and significantly limits the penetration of systemically administered chemotherapeutic agents into the tumor core. As a result, therapeutic agents often fail to reach effective concentrations within the tumor, contributing to treatment resistance and poor clinical outcomes. To overcome these delivery barriers, US-mediated drug delivery has emerged as a promising non-invasive strategy to enhance tissue permeability and drug uptake. US is commonly used in combination with gas-filled contrast agents such as microbubbles, which oscillate and cavitate under acoustic stimulation which can temporarily increase cell membrane permeability and enhance drug delivery into tissues and cells (Bouakaz and Escoffre, 2024). However, microbubbles typically range from 1-10 μm in diameter and are therefore largely confined to the vascular compartment, limiting their ability to penetrate the tumor interstitium (Liu et al., 2025). Nanobubbles (NB), typically smaller than 500 nm, have been proposed as an alternative contrast agent and drug delivery platform because their smaller size allows them to extravasate from tumor vasculature and penetrate deeper into tumor tissue (Chen et al., 2025, Exner and Kolios, 2021). In addition, NBs can be internalized by tumor cells via endocytosis, enabling intracellular delivery strategies and potentially altering intracellular signaling and metabolic pathways (Perera et al., 2022). While NBs are increasingly investigated as both imaging agents and therapeutic carriers, the biological effects of NB internalization and subsequent US exposure on tumor cell behavior remain incompletely understood.

It is particularly important in PDAC, where tumor cells exhibit significant metabolic reprogramming. Many PDAC tumors, especially those harboring KRAS mutations, display altered glucose metabolism, increased glutamine utilization, and changes in mitochondrial function that support survival under hypoxic and nutrient-limited conditions (Li et al, 2019). Because US-induced mechanical stress and NB internalization may influence cellular metabolism, proliferation, and stress signaling pathways, understanding these interactions is important for optimizing US-mediated therapeutic strategies and potentially tailoring treatment protocols to specific tumor phenotypes (Przystupski and Ussowicz, 2022).

In addition to NB internalization, the timing of US exposure relative to NB administration may play a critical role in treatment outcome. Differences in spatial localization of cavitation could lead to distinct biological effects highlighting the importance of systematically investigating US timing in relation to NB delivery. Therefore, the aim of this study is to investigate how NB exposure and the timing of US application influence proliferation, metabolism, and stress responses in pancreatic cancer cells. For this purpose, two pancreatic cancer cell lines are used to represent biological heterogeneity in PDAC: PANC-1 cells, which carry a KRAS mutation and BxPC-3 cells, which are KRAS wild-type. Immediate US exposure after NB addition is compared with delayed US exposure after a one-hour NB incubation and a washing step, allowing assessment of the effects of extracellular versus cell-associated NBs to improve the understanding of how NB localization and US timing influence pancreatic cancer cell behavior and to provide insight into optimizing US-mediated therapeutic strategies for pancreatic cancer.

## Material and Methods

### 1. NB Preparation and Incubation

PGG NBs were produced based on previously described protocols (deLeon, 2019). A lipid mixture composed of DBPC (60.1 mg), DPPE (20 mg), DPPA (10 mg), and DSPE-mPEG2k (10 mg) was made. Propylene glycol (1 mL) was added, and the mixture was heated to 80 °C with sonication until fully dissolved. A pre-heated mixture of glycerol (1 mL) and PBS (8 mL) was then introduced, followed by sonication at room temperature for 10 minutes. The resulting 10 mL lipid suspension was aliquoted and stored at 4 °C for up to two weeks.

For NB formation, 1 mL of lipid solution was transferred into a 3 mL vial. Air was removed using a 25 mL syringe, and the headspace was filled with octafluoropropane (C_3_F_8_) gas. The vial was shaken for 45 seconds using a VialMix (Bristol-Myers Squibb Medical Imaging, MA). The vial was inverted and centrifuged at 550 rpm for 5 minutes to collect NBs at the bottom. Approximately 400 µL of NB suspension was withdrawn using a modified 25G needle.

NB size and concentration were analyzed using the Archimedes resonant mass spectrometry system (Malvern Panalytical). The two time points (t = 0 vs. 1 h) were selected to distinguish between extracellular and intracellular nanobubble localization. Immediate ultrasound exposure (t = 0) primarily reflects cavitation effects from extracellular or membrane-associated NBs, whereas the 1-hour incubation permits cellular uptake and vesicular trafficking, enabling assessment of how intracellular localization influences ultrasound responsiveness.

For the 1-hour condition, a washing step (centrifugation at 200g for 5 minutes) was included to remove non-internalized NBs, ensuring that the observed ultrasound response arises predominantly from cell-associated or internalized nanobubbles, rather than free extracellular particles.

### 2. Cell Culture

Human pancreatic adenocarcinoma cell lines PANC-1 and BxPC-3 were used. PANC-1 cells (ATCC CRL-1469) were cultured in DMEM (Gibco™, Thermo Fisher Scientific, Milano, cat# LS11965092) supplemented with 10% (v/v) FCS (Gibco™, Thermo Fisher Scientific, Milano, cat# 10437-036). Cells were maintained at 37 °C in a humidified incubator with 5% CO_2_ and passaged at a 1:5 ratio using 0.025% Trypsin-EDTA (Thermo Fisher Scientific, cat# 15090046).

BxPC-3 cells (ATCC CRL-1687) were maintained in RPMI 1640 (Gibco™, Thermo Fisher Scientific, Milano, cat# 11875093) supplemented with 10% (v/v) FCS (Gibco™, Thermo Fisher Scientific, Milano, cat# 10437-036) under identical incubation conditions.

### 3. Ultrasound exposure and passive cavitation detection

Ultrasound experiments were performed in a water tank equipped with a 3D positioning system reported previously (Khan et al, 2025). A 1 MHz piezoelectric transducer was used for insonation, and a PVDF passive cavitation detection transducer was positioned orthogonally to record acoustic emissions. Ultrasound settings were 770 kPa peak negative pressure, 1% duty cycle, 500 Hz pulse repetition frequency, and 1 minute treatment duration. Sample containers were made from mylar film. Signal processing was done using MATLAB. Time domain signals were converted in power spectra and normalized with references (cells only) for both cell types. Normalized Power spectral density has been analyzed.

### 4. Cell viability

Viability was determined using an automated Vi-CELL cell viability analyzer (Beckman Coulter), which operates based on Trypan Blue exclusion. After treatment, PANC-1 cells were harvested and gently resuspended to generate a uniform single-cell suspension while minimizing cell aggregation. A portion of each sample was transferred into the Vi-CELL sample cup for analysis. Measurements were conducted immediately following sample preparation, and any necessary dilution factors were incorporated using the instrument’s software.

### 5. MTT Assay

Cellular metabolic activity was assessed using MTT (Sigma #475989) after the USNB treatments. PANC-1 and BxPC-3 cells (10^4^) were seeded into 96-well plates allowed to adhere overnight at 37 °C in a humidified incubator with 5% CO_2_. MTT reagent was added to each well at a final concentration of 0.5 mg/mL, and plates were incubated for 4 hours at 37 °C to allow intracellular conversion of MTT to formazan. After incubation, the medium containing excess MTT was carefully removed, and the resulting formazan crystals were dissolved in dimethyl sulfoxide. Plates were gently shaken to ensure complete dissolution. Absorbance was measured using a microplate reader at 570 nm, with background subtraction 690 nm. Blank wells containing medium and MTT, but no cells were used for background correction. The absorbance values were normalized to untreated control cells within each experiment, with controls set to 100%. All conditions were analyzed in 4 technical replicates, and three independent biological experiments were performed.

### 6. Ki67 Staining

Cell proliferation in PANC-1 and BxPC-3 cells was evaluated using Ki-67 immunofluorescence (Abcam, #231172) in combination with nuclear staining using DAPI (ThermoFisher, #D1306). Poly-D-lysine-coated 15 mm glass coverslips were placed into 12-well plates, and 2 × 10^5^ cells were seeded per well in 500 µL of complete culture medium. After allowing cells to adhere, they were washed with PBS, fixed with 4% (v/v) paraformaldehyde, and permeabilized using 0.1% (v/v) Triton X-100 in PBS. To reduce non-specific binding, cells were blocked for 1 hour at room temperature in a solution containing 5% goat serum and 3% BSA. Samples were then incubated overnight at 4 °C with anti-Ki-67 primary antibody (1:250 dilution). The following day, cells were incubated for 1 hour at room temperature with goat anti-rabbit Alexa Fluor 555 secondary antibody (Invitrogen #A-21428) (1:500) along with DAPI (1:5000).

Coverslips were mounted using DAKO mounting medium. Fluorescence images were captured using an Olympus BX50 microscope, with five distinct non-overlapping fields acquired per coverslip. Quantification of DAPI-positive nuclei and Ki-67-positive cells was performed using Fiji/ImageJ software. For each biological replicate, approximately 150 cells were analyzed per field across five non-overlapping regions (∼750 cells total per sample). The counts from these regions were averaged to obtain a single value per replicate. Data were normalized to the untreated control within each experiment (set to 100%). Statistical analysis was carried out using GraphPad Prism 10.

### 7. Protein Extraction and Quantification

Following the treatments (1 × 10^6^) cells were seeded on 6-well plates, after overnight incubation, lysates were prepared using R&D Systems kit (ARY018). Protein concentration was determined using Pierce™ BCA Kit (Thermo Scientific #: 23227).

### 8. Cell Stress Protein Array Analysis

Protein profiling was performed using Proteome Profiler Human Cell Stress Array Kit (R&D# ARY018). 150 µg protein was incubated with detection antibodies and applied to membranes. After overnight incubation, membranes were developed and analyzed using Bio-Rad ChemiDoc and Quick Spots software.

### 9. Statistical Analysis

Data were analyzed using GraphPad Prism 8. Mean ± SD is reported. Two-way ANOVA with Tukey’s or Sidak’s test was used. Significance levels: (*) P < 0.05, (**) P < 0.01, (***) P < 0.001, (****) P < 0.0001.

## Results

### Reduced cavitation activity when US is applied after NB incubation and washing

To determine whether the timing of US relative to NB exposure affects cavitation activity, passive cavitation detection was performed when US was applied either immediately after NB addition (t = 0) or after a 1-hour incubation followed by washing to remove extracellular NBs.

In both cell lines a strong broadband and harmonic acoustic emissions were detected when US was applied immediately after NB addition. In contrast, when cells were incubated with NBs for 1 hour that were washed, and then exposed to US, the acoustic emissions were markedly reduced but it was still there. This reduction in signal intensity suggests that freely suspended extracellular NBs contribute significantly to cavitation activity, whereas after washing, only cell-associated or internalized NBs remain they are still acoustically active with reduced cavitation signals (Figure 1 a, b and Supplemental video 1a-d).

**Figure 1.**
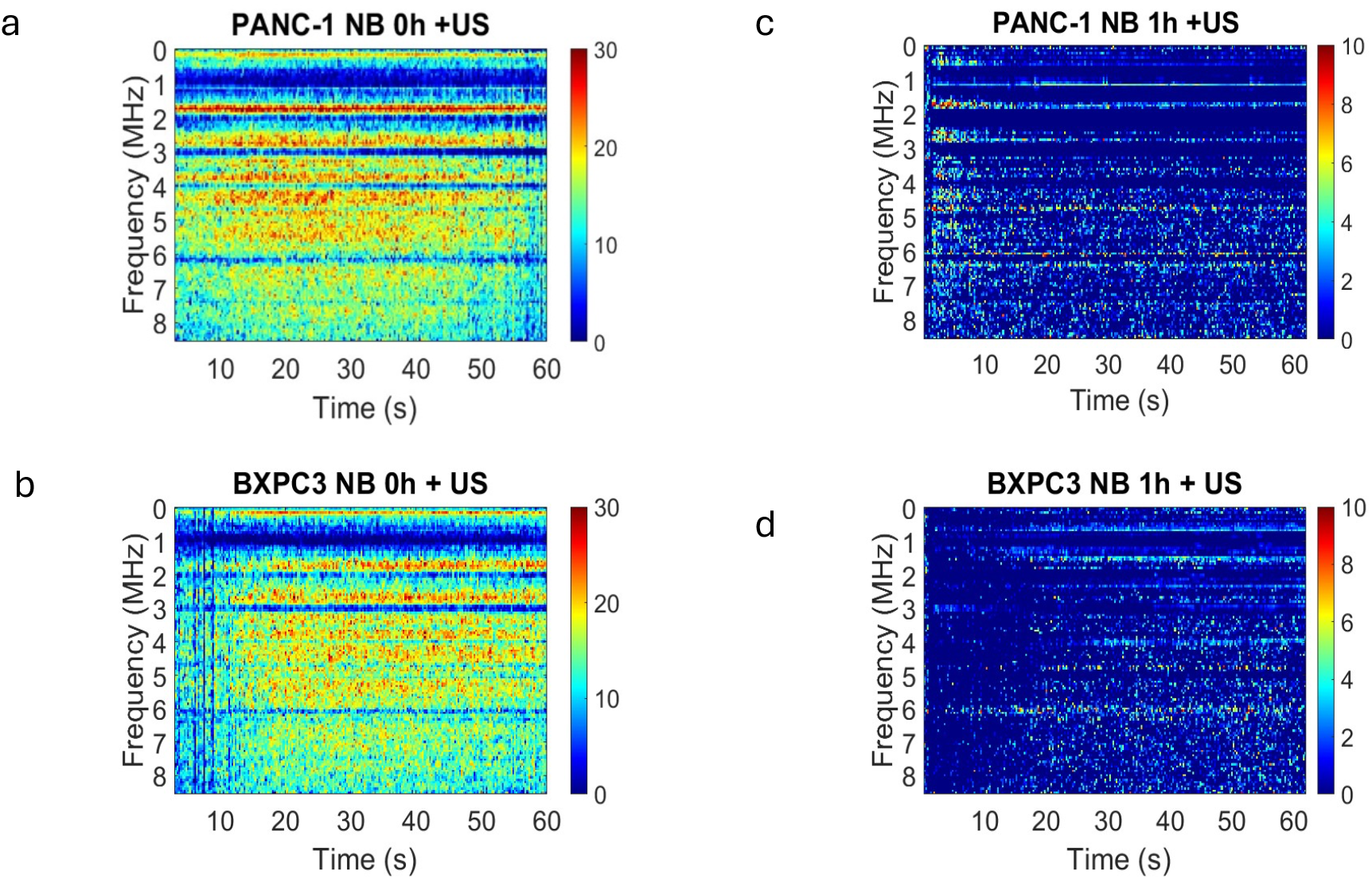
Passive cavitation detection during US exposure under immediate and delayed NB conditions. Passive cavitation detection spectrograms recorded during US exposure of pancreatic cancer cells treated with NBs. US was applied either immediately after NB addition (t = 0, extracellular NBs present) or after a 1-hour incubation followed by washing to remove extracellular NBs (1 h wash). Spectrograms show acoustic emissions for PANC-1 and BxPC-3 cells under both conditions. Immediate US exposure produced stronger broadband and harmonic emissions, whereas delayed US after washing resulted in reduced acoustic signal intensity, indicating reduced cavitation activity when extracellular NBs were removed and primarily cell-associated NBs remained (Supplemental Video1)

**Supplemental Figure 1 (Video):**
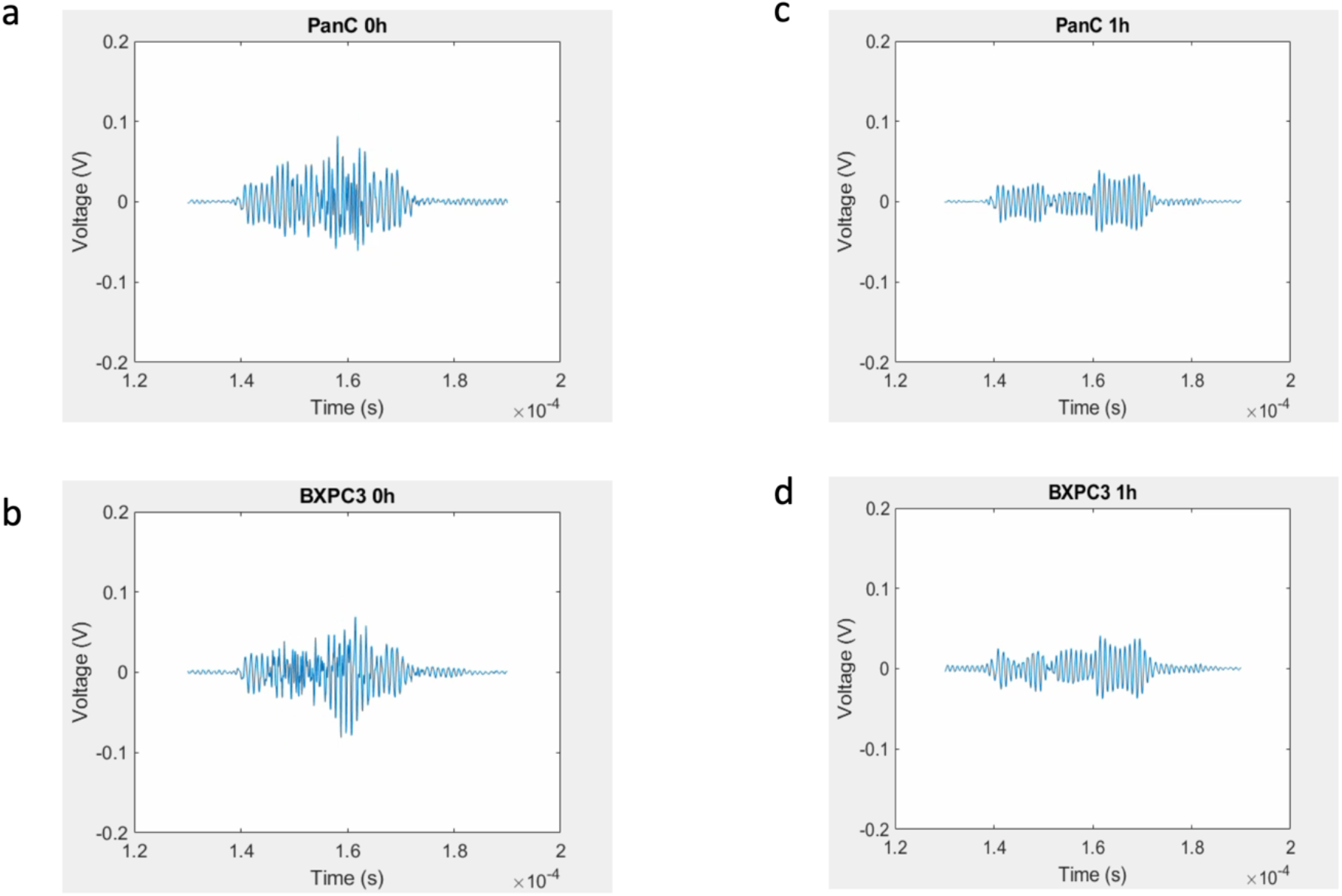
Immediate US exposure produced stronger broadband and harmonic emissions, whereas delayed US after washing resulted in reduced acoustic signal intensity, indicating reduced cavitation activity when extracellular NBs were removed and primarily cell-associated NBs remained.

These results confirm that the timing of US exposure relative to NB incubation significantly affects cavitation behavior, likely due to the presence or absence of extracellular NBs at the time of insonation.

### Immediate US exposure in the presence of extracellular NBs reduces cell viability more than delayed US exposure after washing

Having observed differences in cavitation activity between immediate and delayed US exposure, we next asked whether these differences translated into differences in cell viability. Cell viability was measured in PANC-1 and BxPC-3 cells using trypan blue exclusion at 1 hour (Figure 2a) and 24 hours after treatment (Figure 2b). At 1 hour after treatment, viability differences between treatment groups were minimal, indicating that treatments did not cause immediate cell death. Since these cells have doubling time more than 24 hours we wanted to observe the viability at 24 hours after the treatment. Viability was indeed decreased in cells treated with US and NBs compared with untreated and US-only controls at 24 hours. Importantly, the greatest reduction in viability was observed when US was applied immediately after NB addition (t = 0), when extracellular NBs were still present. When US was applied after a 1-hour incubation and washing step, the reduction in viability was smaller suggesting that cavitation generated by extracellular NBs produces stronger cytotoxic effects than US exposure applied after removal of extracellular NBs. Also, the delayed reduction in viability indicated that treatment-induced damage may lead to delayed cell death rather than immediate apoptosis.

**Figure 2.**
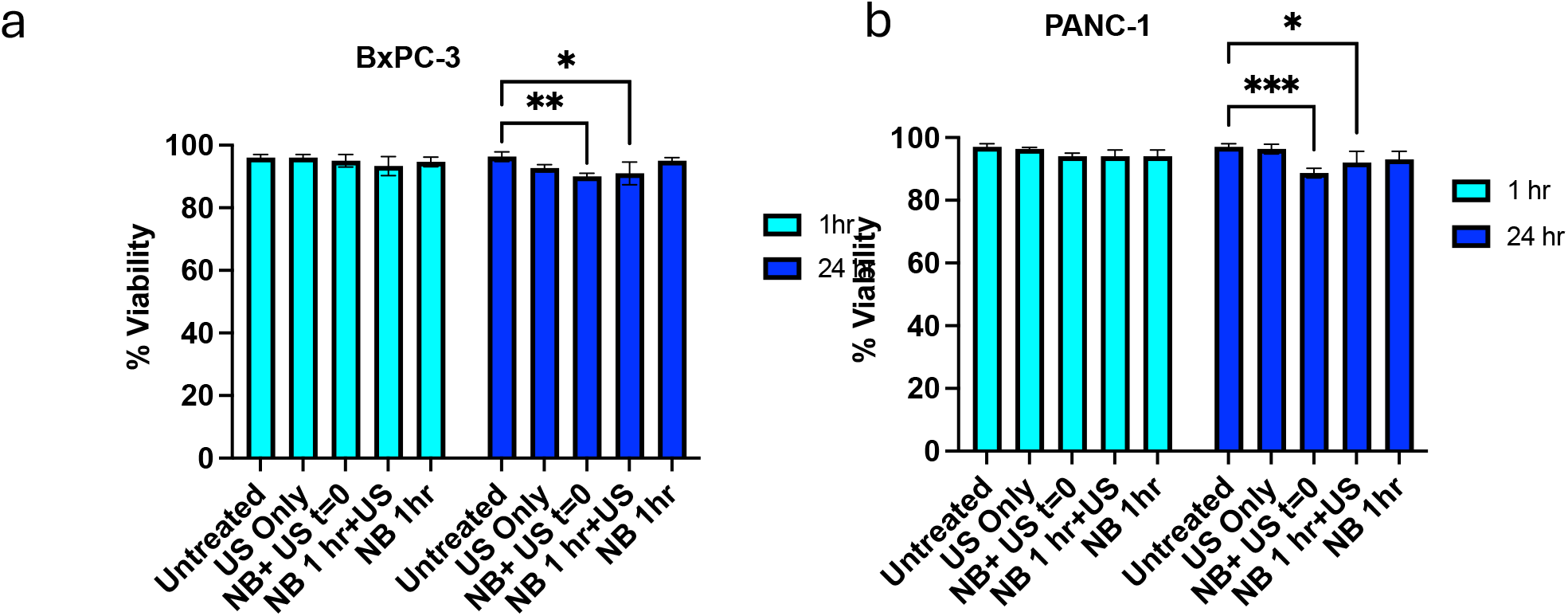
Cell viability following US and NB treatments. Cell viability measured using trypan blue exclusion in BxPC-3 and PANC-1 cells at 1 hour and 24 hours after treatment. Treatment groups included untreated control, US only, US applied immediately after NB addition (NB + US t = 0), US applied after 1-hour NB incubation followed by washing (NB 1 h + US), and NBs only (NB 1 h). Immediate US exposure in the presence of extracellular NBs resulted in a greater reduction in viability compared with delayed US exposure after washing. Viability differences were more pronounced at 24 hours than at 1 hour, indicating delayed treatment effects. Data are presented as percent viability relative to untreated control. Statistical significance is indicated, n=8, (*) P < 0.05, (**) P < 0.01.

### US and NB treatments reduce proliferative activity in a timing-dependent manner

Since viability differences were more pronounced after 24 hours, and both BxPC-3 and PANC-1 cells’ doubling time is more than 24 hours, we next investigated whether US and NB treatments affected cell proliferation. Cell proliferation was evaluated using Ki-67 immunofluorescence staining in BxPC-3 and PANC-1 cells (Figures 3 and 4). In both cell lines, the percentage of Ki-67-positive proliferating cells decreased following treatments involving US and NBs compared with untreated controls. Quantification of Ki67-positive cells showed that untreated cells exhibited approximately 40% proliferating cells. US-only treatment reduced the percentage of proliferating cells to approximately 30%. The combination of US and NBs resulted in a further reduction in proliferative activity, with the US+NBs 0T group showing approximately 25% proliferating cells and the US+NBs 1T group showing the lowest proliferation at approximately 20%. In contrast, the NBs-only group showed increased proliferative activity compared to untreated cells, with approximately 50% Ki67-positive cells. Overall, these results indicate that NBs alone increased proliferative activity, whereas the combination of NBs and US reduced proliferation in BxPC-3 cells, with the greatest reduction observed in the US+NBs 1T group. In PANC-1 cells, untreated cells exhibited a higher proliferative fraction, with approximately 60–70% Ki67-positive cells. US-only treatment reduced proliferation to approximately 55%. The US+NBs 0T group showed a further reduction to approximately 45% proliferating cells, while the US+NBs 1T group showed the greatest reduction, with approximately 35% Ki67-positive cells. In contrast, the NBs-only group showed increased proliferation compared to untreated cells, with approximately 75% proliferating cells. Overall, these results indicate that the combination of NBs and US reduced proliferative activity in both cell lines, with a more pronounced reduction observed in PANC-1 cells compared to BxPC-3 cells.

**Figure 3.**
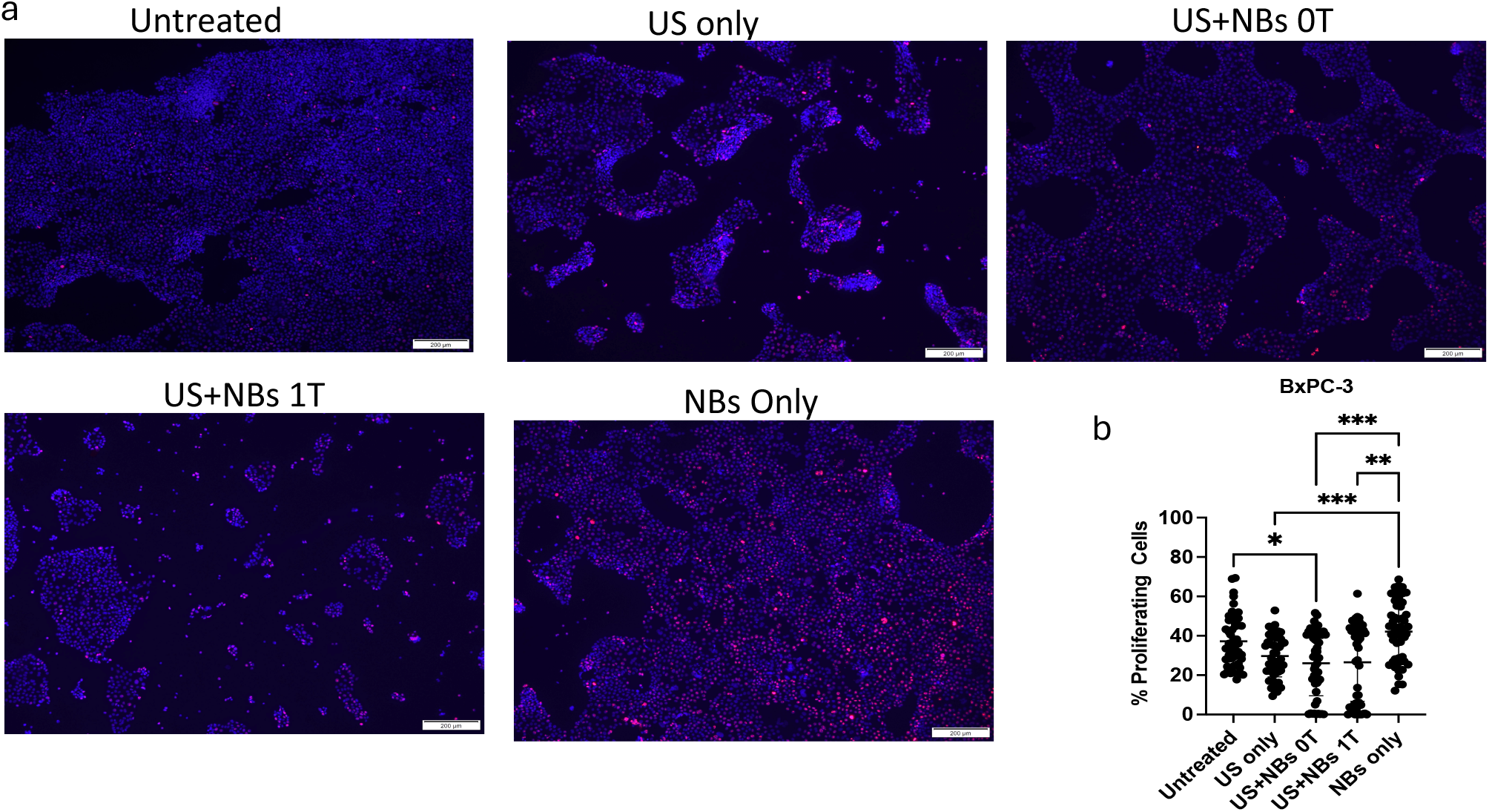
Ki-67 proliferation analysis in BxPC-3 cells following US and NB treatment. (a) Representative immunofluorescence images of BxPC-3 cells stained for Ki-67 (proliferation marker) and nuclei (DAPI) under different treatment conditions: untreated, US only, US applied immediately after NB addition (NB + US 0T), US applied after 1-hour NB incubation and washing (NB + US 1T), and NBs only. (b) Quantification of the percentage of Ki-67 positive proliferating cells for each treatment group. US combined with NBs significantly reduced proliferative activity compared with controls, with the greatest reduction observed when US was applied immediately after NB addition. Data are presented as mean ± SD with statistical significance indicated. (*) P < 0.05, (**) P < 0.01, (***) P < 0.001.

**Figure 4.**
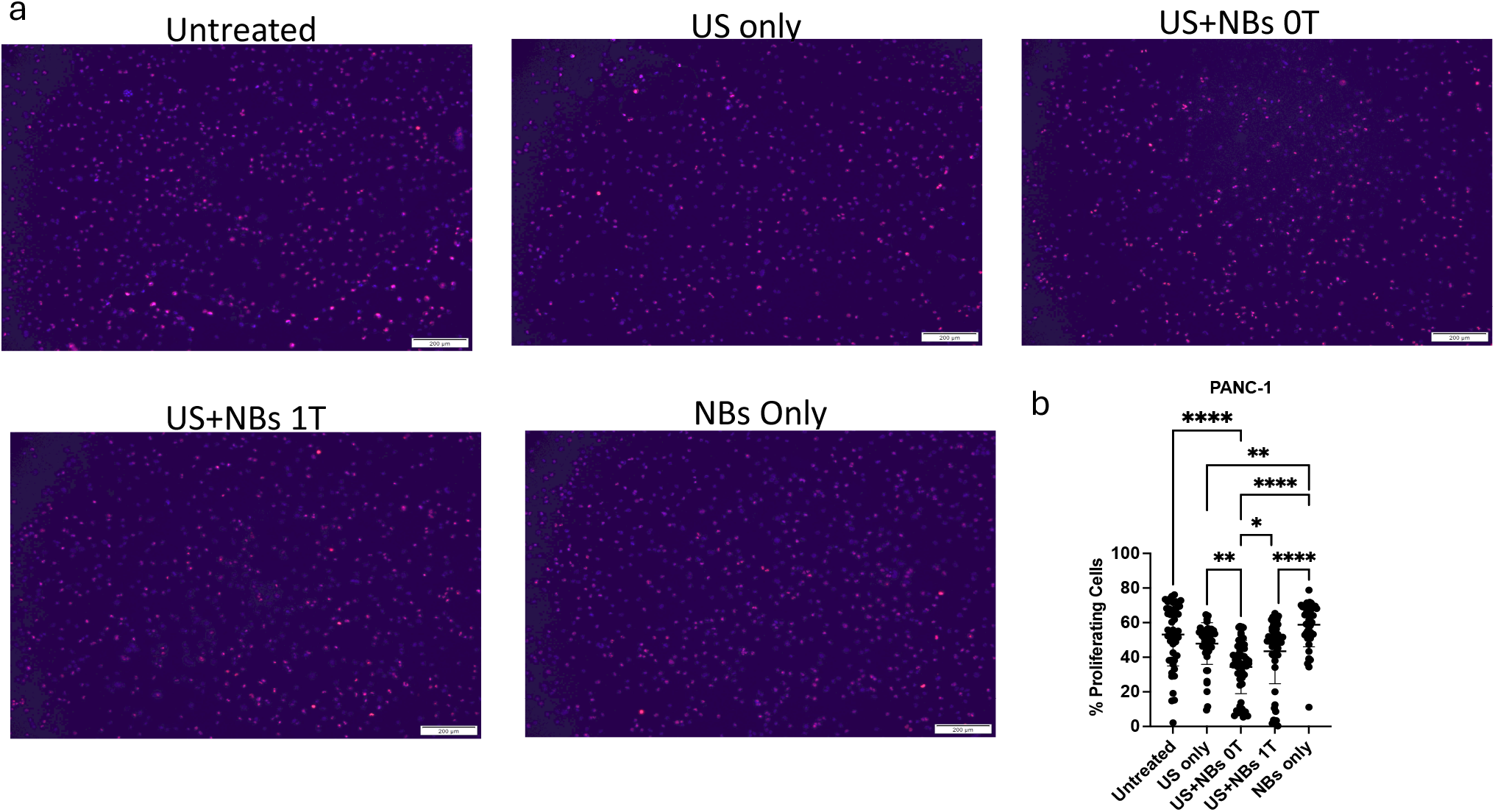
Ki-67 proliferation analysis in PANC-1 cells following US and NB treatment. (a) Representative immunofluorescence images of PANC-1 cells stained for Ki-67 and DAPI under different treatment conditions: untreated, US only, US applied immediately after NB addition, US applied after 1-hour NB incubation and washing, and NBs only. (b) Quantification of the percentage of proliferating cells for each treatment group. Both US and US combined with NBs reduced proliferative activity compared with untreated controls, with timing-dependent differences observed between immediate and delayed US exposure. Data are presented as mean ± SD with (*) P < 0.05, (**) P < 0.01, (***) P < 0.001, (****) P < 0.0001.

In BxPC-3 cells, US alone produced a modest reduction in proliferation, whereas US combined with NBs resulted in a greater reduction (Figure 3 a,b). The largest decrease in proliferating cells was observed when US was applied immediately after NB addition. Similarly, in PANC-1 cells, immediate US exposure in the presence of extracellular NBs resulted in one of the strongest reductions in proliferative activity, while delayed US exposure after washing also reduced proliferation but generally to a lesser extent (Figure 4 a,b). proliferation is more sensitive to treatment timing than immediate viability suggesting that US-NB exposure may induce cell cycle arrest or delayed growth inhibition even in cells that initially remain viable. Moreover, US only treatment produced a modest reduction in proliferative activity in both cell lines, decreasing Ki-67-positive cells from 40% to 30% in BxPC-3 and from 65% to 55% in PANC-1, indicating that US can also influence cell cycle dynamics even in the absence of NBs.

### US and NB treatments influence metabolic activity and reveal intrinsic metabolic differences between pancreatic cancer cell lines

Since proliferation and viability were affected by treatment timing, we next asked whether US and NB treatments also influenced MTT uptake and cellular metabolism. MTT signal was measured at 24, 48, and 72 hours post-treatment to assess treatment-associated changes in cellular metabolic activity in BxPC-3 and PANC-1 cells following NB and US exposure. In BxPC-3 cells, untreated cells showed a gradual increase in MTT signal over time. US-only treatment produced a similar trend with a modest increase over time. NB treatment alone (NB 1 hr) resulted in a higher MTT signal compared to untreated controls at all time points, indicating increased metabolic activity. In contrast, NB+US t=0 treatment resulted in reduced MTT signal compared to untreated cells at 24, 48, and 72 hours. The NB 1 hr + US group showed intermediate MTT values between NB alone and NB+US t=0 groups, with MTT signal increasing over time but remaining lower than the NB-only group (Figure 5a).

**Figure 5.**
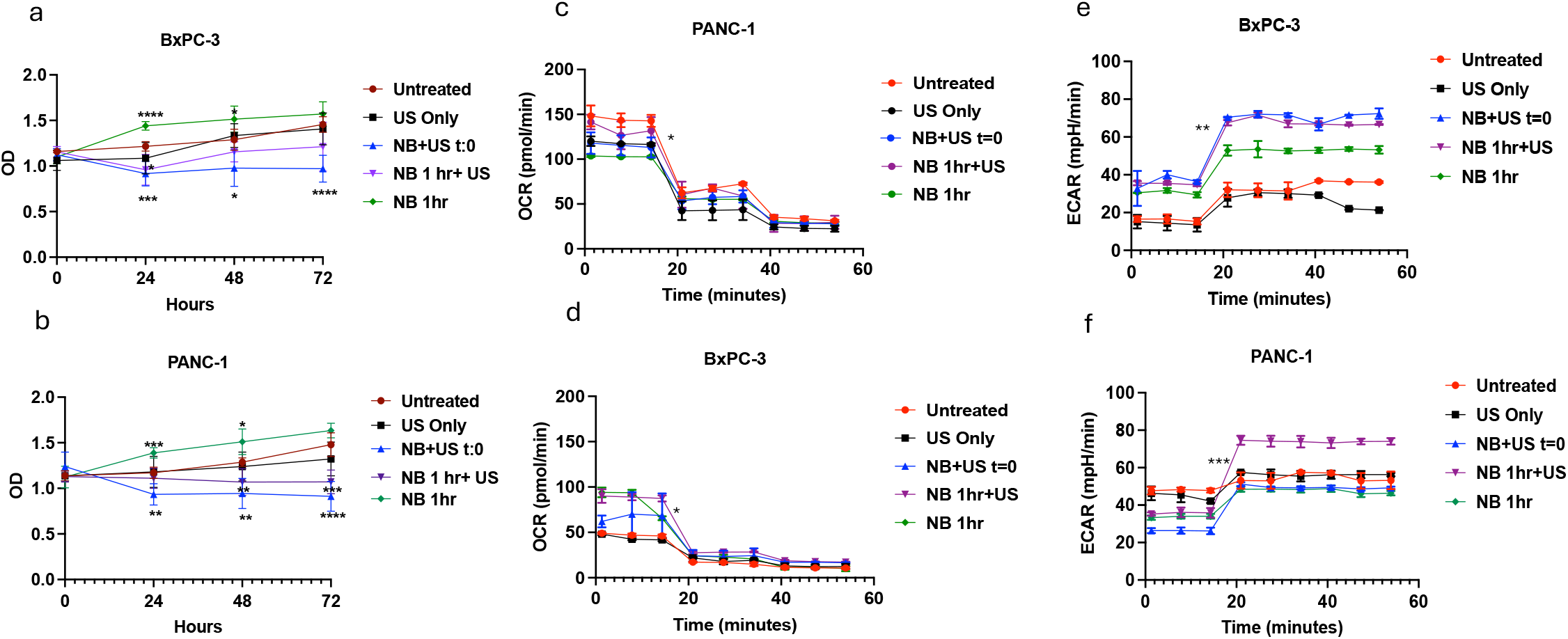
Metabolic activity and Seahorse bioenergetic analysis following US and NB treatment. (a-b) Optical density measurements over 72 hours representing metabolic activity measured using the MTT assay in BxPC-3 (a) and PANC-1 (b) cells following treatment. (c-d) Extracellular acidification rate (ECAR) measurements over time for BxPC-3 (c) and PANC-1 (d) cells, indicating glycolytic activity. (e-f) Oxygen consumption rate (OCR) measurements over time for BxPC-3 (e) and PANC-1 (f) cells, indicating mitochondrial respiration. Treatment groups included untreated control, US only, US applied immediately after NB addition, US applied after 1-hour NB incubation and washing, and NBs only. The data demonstrate intrinsic metabolic differences between cell lines and timing-dependent metabolic responses to US-NB treatments. Data were analyzed using GraphPad Prism 8. Mean ± SD is reported with two-way ANOVA with Tukey’s (*) P < 0.05, (**) P < 0.01, (***) P < 0.001, (****) P < 0.0001.

**Figure 6.**
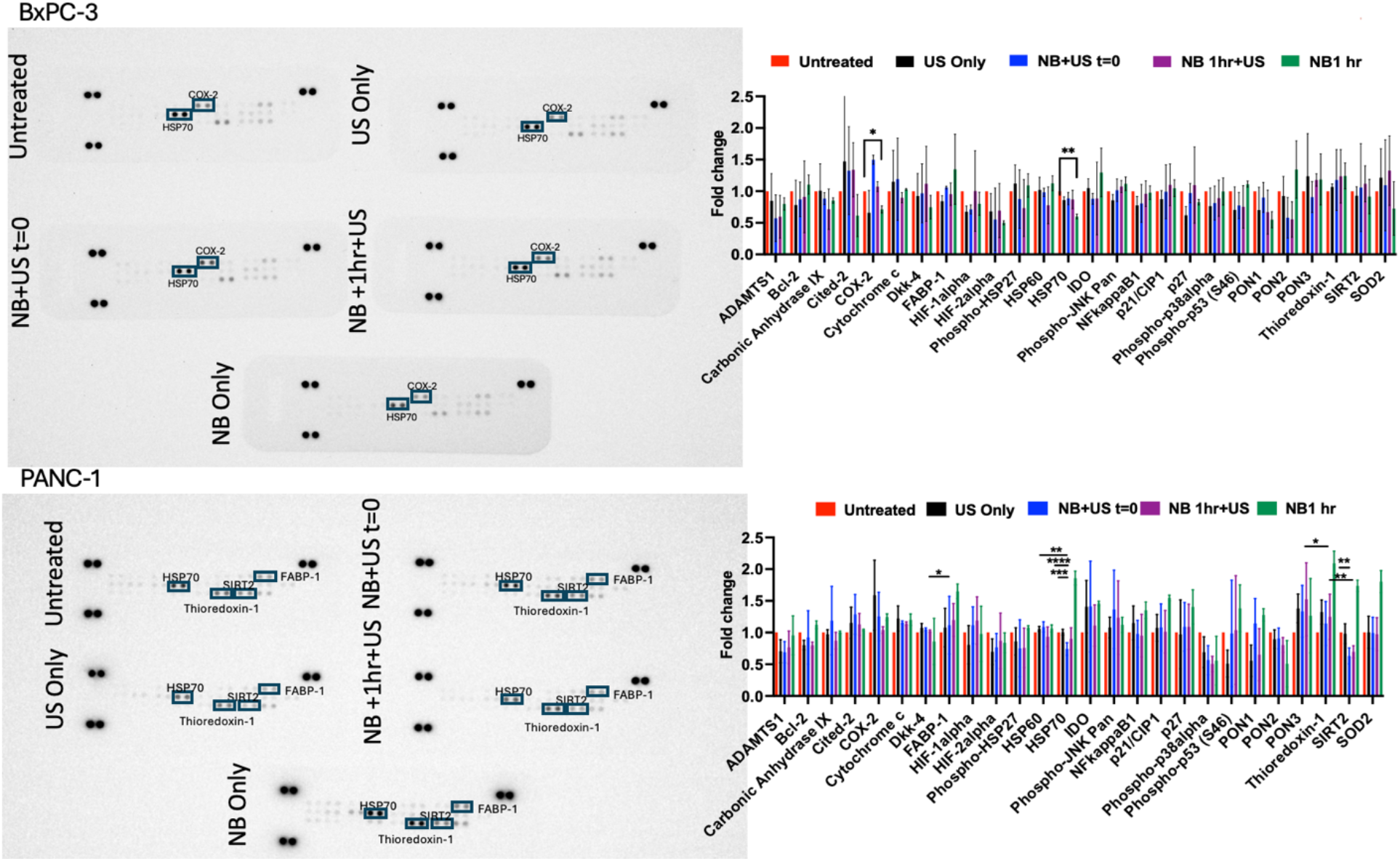
Human cell stress protein array analysis following US and NB treatments. Representative human cell stress array membranes and quantification of relative protein expression in BxPC-3 and PANC-1 cells across treatment conditions. The array included proteins associated with oxidative stress, hypoxia response, apoptosis, and stress signaling pathways. Bar graphs show relative expression or fold change compared with untreated controls. PANC-1 cells exhibited broader stress protein expression changes in proteins HSP70, Thioredoxin-1, SIRT-2 and FABP-1 compared with BxPC-3 cells that had changes in expression of Cox-2 and HSP-70, n=3, Mean ± SD is reported with two-way ANOVA with Tukey’s (*) P < 0.05, (**) P < 0.01, (***) P < 0.001, (****) P < 0.0001.

In PANC-1 cells, untreated cells again showed a gradual increase in MTT signal over time. NB treatment alone (NB 1 hr) resulted in increased MTT signal compared to untreated controls at all time points. In contrast, NB+US t=0 treatment resulted in a marked decrease in MTT signal compared to untreated cells at 24, 48, and 72 hours. The NB 1 hr + US group also showed reduced MTT signal compared to untreated cells, although the reduction was less pronounced than in the NB+US t=0 group. US-only treatment showed a gradual increase in MTT signal over time but remained lower than untreated controls at later time points (Figure 5b).

Furthermore, to assess metabolic changes in BxPC-3 and PANC-1 pancreatic cancer cells following NB and US treatment seahorse extracellular flux analysis was used by measuring oxygen consumption rate (OCR) and extracellular acidification rate (ECAR). Oligomycin was injected to inhibit ATP synthase and determine ATP linked respiration and compensatory glycolysis, followed by rotenone to inhibit mitochondrial respiration and determine non-mitochondrial oxygen consumption. Basal OCR measurements showed that in BxPC-3 cells, NB-treated groups exhibited moderately higher oxygen consumption compared to untreated and US-only groups. Basal OCR values ranged from approximately 45-50 pmol/min in untreated cells to approximately 90-95 pmol/min in the NB 1 hr group, indicating increased mitochondrial respiratory activity following NB exposure. In contrast, PANC-1 cells showed overall higher basal OCR values compared to BxPC-3 cells, ranging from approximately 105-150 pmol/min across treatment groups, with untreated cells exhibiting the highest basal respiration and treated groups generally showing slightly reduced basal OCR (Figure 5c).

After rotenone injection, OCR decreased further to low residual levels consistent with non-mitochondrial respiration, reaching approximately 12-18 pmol/min in BxPC-3 cells and approximately 25-35 pmol/min in PANC-1 cells (Figure 5d).

ECAR increased in both cell lines following oligomycin injection, indicating compensatory glycolysis when mitochondrial ATP synthesis was inhibited. In BxPC-3 cells, the increase in ECAR was most pronounced in the NB+US t=0 and NB 1 hr + US groups, which showed higher ECAR values compared to untreated and US-only groups, indicating enhanced glycolytic activity following treatment. The NB 1 hr group also showed increased ECAR relative to controls, although to a lesser extent, suggesting that NB treatment, particularly when combined with US, enhances glycolytic activity and overall metabolic responsiveness in BxPC-3 cells (Figure 5e).

In PANC-1 cells, ECAR also increased following oligomycin injection in all treatment groups; however, the increase was most substantial in the NB 1 hr + US group, which exhibited the highest ECAR values compared to all other groups. Other treatment groups showed moderate ECAR increases compared to untreated controls. Compared to BxPC-3 cells, PANC-1 cells demonstrated a stronger glycolytic response relative to mitochondrial respiration following treatment, indicating a shift toward glycolytic metabolism, particularly following NB 1 hr + US treatment indicating a more evident shift toward glycolytic metabolism in PANC-1 cells following treatment (Figure 5f).

### US and NB treatments activate cellular stress signaling pathways, particularly in PANC-1 cells

To investigate whether the observed changes in metabolism and proliferation were associated with cellular stress responses, a human cell stress protein array was performed following US and NB treatments (Figure X). Differential expression of several stress-related proteins was observed in both BxPC-3 and PANC-1 cells, with more pronounced changes observed in PANC-1 cells.

In BxPC-3 cells, relatively modest changes in stress protein expression were observed across treatment groups. HSP70 expression showed a slight increase in the NB+US t=0 group (1.4-fold) compared to untreated cells, while NB only treatment also showed a mild increase (1.3-fold). COX-2 expression showed small variations across treatments, generally ranging between 0.8 and 1.3-fold. Overall, stress protein changes in BxPC-3 cells were relatively modest, indicating a limited stress response compared to PANC-1 cells.

In contrast, PANC-1 cells exhibited a broader and more pronounced stress response across multiple proteins. HSP70 expression increased in several treatment groups, with the highest expression observed in the NB 1 hr + US group (∼2.0-fold), while the NB only group also showed increased HSP70 expression (1.6-fold) compared to untreated cells. Thioredoxin-1 expression increased in both NB only and NB+US treatment groups, reaching approximately 1.8-fold relative to untreated cells. SIRT2 expression also increased following NB treatments, with values approximately 1.7-fold in NB-containing groups compared to untreated cells. FABP-1 expression was almost 2-fold increase in NB only group.

In sum, NB treatment alone induced changes in several stress-related proteins, and the combination of NBs and US further enhanced the stress response, particularly in PANC-1 cells. Therefore, NBs contribute significantly to the cellular stress response, and that US in combination with NBs amplifies stress signaling pathways, particularly those associated with oxidative stress, heat shock response, and metabolic stress signaling.

## Discussion

PDAC remains one of the most challenging malignancies to treat due to its aggressive biology, late diagnosis, and the presence of a dense desmoplastic stroma that limits drug delivery and therapeutic efficacy. US-mediated drug delivery using acoustic contrast agents such as microbubbles and NBs has emerged as a promising strategy to enhance drug delivery by increasing tissue permeability and facilitating drug uptake through cavitation-induced sonoporation. However, while NBs are increasingly investigated as delivery vehicles and imaging agents, their direct biological effects on tumor cells remain incompletely understood. In this study, we investigated how NB exposure, with and without US and under different timing conditions, affects viability, proliferation, metabolism, and stress signaling in pancreatic cancer cells.

Our passive cavitation results demonstrated that acoustic emissions were significantly stronger when US was applied immediately after NB addition compared with US applied after a 1-hour incubation and washing step indicating that freely suspended extracellular NBs contribute significantly to cavitation activity, whereas after washing, cavitation activity is reduced due to the removal of extracellular bubbles and the presence of only cell-associated or internalized NBs. These differences in cavitation activity were reflected in the biological outcomes observed in the viability and proliferation experiments as reduced viability and proliferation. Although immediate cell death was minimal shortly after treatment, viability decreased significantly 24 hours after US exposure in the USNB treated groups most notably when NBs are mostly extracellular suggesting that mechanical stress generated during cavitation may induce delayed apoptosis or growth arrest. Such delayed cytotoxic effects have been reported in US-mediated sonoporation studies, where membrane disruption, calcium influx, oxidative stress, and mitochondrial dysfunction lead to delayed cell death or proliferation arrest rather than immediate cell lysis (Sharma, D et al., 2019; McNabb et al, 2020).

Along with the reduced viability in USNB treated groups, proliferative activity measured by Ki-67 staining was also reduced in both cell lines suggesting that USNB exposure may induce cell cycle arrest or stress-mediated growth inhibition rather than acute cytotoxicity. The metabolic assays provided further insight into the cellular responses to NBs and US. The MTT assay showed time-dependent changes in metabolic activity over 72 hours, with immediate US plus NBs reducing metabolic activity, while NBs incubated for 1 hour without US sometimes increased MTT signal. Because these metabolic changes were not fully consistent with Ki-67 proliferation data, the MTT response likely reflects changes in cellular redox state or mitochondrial activity rather than simply changes in cell number. Since MTT reduction depends on cellular NAD(P)H dependent oxidoreductase activity, changes in MTT signal may reflect altered metabolic state, mitochondrial function, or oxidative stress rather than proliferation alone (Ghasemi et al, 2021).

Seahorse metabolic flux analysis further demonstrated differences in metabolic responses between the two pancreatic cancer cell lines. PANC-1 cells, which harbor KRAS mutations, showed a more glycolytic phenotype compared with BxPC-3 cells, consistent with previous reports showing that KRAS mutations drive metabolic reprogramming toward glycolysis and altered mitochondrial metabolism in pancreatic cancer (Zhang et al, 2025). Following oligomycin injection, OCR decreased and ECAR increased in both cell lines, indicating a compensatory shift toward glycolysis when mitochondrial respiration was inhibited. Glycolytic response was more pronounced in PANC-1 cells, particularly in NB and US treatment groups, suggesting that treatment may enhance glycolytic reliance or metabolic stress adaptation in this cell line. The metabolic findings suggest that US and NB exposure influences cellular metabolism and metabolic balance rather than completely disrupting energy production.

To explore whether US and NB exposure induced cellular stress responses, we performed a human stress protein array. PANC-1 cells showed a broader activation of stress-related proteins compared with BxPC-3 cells, including increased expression of proteins associated with oxidative stress, heat shock response, and metabolic stress signaling such as HSP70, thioredoxin-1, SIRT2, and FABP-1 suggesting that cellular characteristics such as genotype and metabolic phenotype influence the response to USNB treatment, and that more glycolytic PANC-1 cells may be more sensitive to stress signaling pathways triggered by mechanical or metabolic perturbations.

Among the stress proteins analyzed, HSP70 showed an interesting pattern: HSP70 expression was commonly changed in both BxPC-3 and PANC-1 cells upon treatments but the response was different among the cell lines. It was decreased in USNB-treated groups in PANC-1 and increased in NB only group compared with untreated controls, whereas in BxPC-3 cells, it was decreased in Following 1-hour incubation with nanobubbles (NB), differential stress responses were observed between PANC-1 and BxPC-3 cells. HSP70, a key molecular chaperone involved in protein folding, oxidative stress protection, and inhibition of apoptosis, is commonly upregulated in response to cellular stress and is associated with tumor aggressiveness (Giri et al., 2017). In PANC-1 cells, NB treatment alone led to an increase in HSP70 expression, suggesting activation of an adaptive stress response. However, when US was combined with NB, HSP70 expression decreased that may be attributed to US induced mechanical effects, such as cavitation and membrane disruption, which can cause protein denaturation or oxidative stress levels that exceed the protective capacity of HSP70. Additionally, US exposure may shift cells from adaptive stress responses toward apoptotic signaling or cell cycle arrest pathways, consistent with the observed reduction in Ki-67 expression, indicating decreased proliferation even in the absence of immediate effects on cell viability.

In contrast, BxPC-3 cells exhibited a different response profile. Both HSP70 and COX-2 expression were reduced in NB-only groups, with minimal additional changes following US treatment suggesting that NB exposure alone may be sufficient to suppress basal stress-survival and inflammatory signaling in BxPC-3 cells that may be affected by US exposure after 1 hour NB incubation.

To sum up, our results demonstrate that the biological effects of USNB treatment depend on the timing of US exposure relative to NB incubation. Immediate US exposure in the presence of extracellular NBs produces stronger cavitation, greater reductions in viability, and stronger inhibition of proliferation. In contrast, delayed US exposure after washing results in reduced cavitation and smaller effects on viability but still influences proliferation, metabolism, and stress signaling, likely through intracellular or signaling mediated mechanisms rather than direct mechanical damage.

## Conclusion

This study demonstrates that NBs can influence tumor cell behavior, including proliferation, metabolism, and cellular stress responses. NBs alone induced adaptive cellular stress and metabolic responses, whereas the combination of NBs and US generated stronger mechanical and metabolic stress, leading to reduced proliferative activity and decreased viability.

Importantly, the biological effects were strongly dependent on the timing of US exposure relative to NB incubation, highlighting treatment timing as a critical parameter in US-mediated therapeutic strategies. Furthermore, the responses differed between pancreatic cancer cell lines, suggesting that tumor metabolic phenotype and cellular characteristics influence sensitivity to USNB treatment providing an important insight into the biological effects of USNB and support its potential as a strategy for modulating tumor cell metabolism and proliferation in pancreatic cancer therapy.

## Acknowledgements

This research was supported by the Natural Sciences and Engineering Research Council of Canada (NSERC) Discovery Grant (RGPIN 2017-05193) and the National Institutes of Health (NIH) Grant 5R01EB028144. The authors would like to thank Simran Bhaskar, Dhruvi Patel, Elizabeth Berndl, and the Keenan Research Centre for Biomedical Science Core Facilities at St. Michael’s Hospital, Toronto, ON, Canada. The authors also wish to thank Lilia Baev at SPARC BioCentre, Hospital for Sick Children, Toronto, Canada for performing Seahorse.

## References

Bouakaz A, Michel Escoffre J. From concept to early clinical trials: 30 years of microbubble-based US-mediated drug delivery research. Adv Drug Deliv Rev. 2024;206:115199. doi:10.1016/j.addr.2024.115199

Chen, L. E., Nittyacharn, P., & Exner, A. A. (2025). Progress and potential of NBs for US-mediated drug delivery. Expert opinion on drug delivery, 22(7), 1007–1030. 10.1080/17425247.2025.2505044

de Leon, A., Perera, R., Hernandez, C., Cooley, M., Jung, O., Jeganathan, S., Abenojar, E., Fishbein, G., Sojahrood, A. J., Emerson, C. C., Stewart, P. L., Kolios, M. C., & Exner, A. A. (2019). Contrast enhanced US imaging by nature-inspired ultrastable echogenic NBs. Nanoscale, 11(33), 15647– 15658. 10.1039/c9nr04828f

Exner, Agata A., and Michael C. Kolios. 2021. “Bursting Microbubbles: How NB Contrast Agents Can Enable the Future of Medical US Molecular Imaging and Image-guided Therapy.” Current Opinion in Colloid & Interface Science 54: 101463. 10.1016/j.cocis.2021.101463.e

Ghasemi, M., Turnbull, T., Sebastian, S., & Kempson, I. (2021). The MTT Assay: Utility, Limitations, Pitfalls, and Interpretation in Bulk and Single-Cell Analysis. International journal of molecular sciences, 22(23), 12827. 10.3390/ijms222312827

Giri, B., Sethi, V., Modi, S., Garg, B., Banerjee, S., Saluja, A., & Dudeja, V. (2017). “Heat shock protein 70 in pancreatic diseases: Friend or foe”. Journal of surgical oncology, 116(1), 114–122. 10.1002/jso.24653

Khan, M. S., Bulycheva, V., Ferworn, C., Falou, O., Strohm, E. M., Nittyacharn, P., Berndl, E., Karshafian, R., Exner, A. A., & Kolios, M. C. (2025). Evaluating the Effect of Drug Loading on the Acoustic Response of Nanobubbles in Stable and Inertial Cavitation Regimes. IEEE open journal of ultrasonics, ferroelectrics, and frequency control, 5, 190–203. 10.1109/ojuffc.2025.3618629

Liu, D., Ling, Y., Liu, W., Deng, J., & Guo, Y. (2025). US-responsive micro/NBs based intelligent theranostic systems for precision tumor therapy. Journal of nanobiotechnology, 24(1), 83. 10.1186/s12951-025-03959-x

Li, J. T., Wang, Y. P., Yin, M., & Lei, Q. Y. (2019). Metabolism remodeling in pancreatic ductal adenocarcinoma. Cell stress, 3(12), 361–368. 10.15698/cst2019.12.205

McNabb, E., Al-Mahrouki, A., Law, N., McKay, S., Tarapacki, C., Hussein, F., & Czarnota, G. J. (2020). Ultrasound-stimulated microbubble radiation enhancement of tumors: Single-dose and fractionated treatment evaluation. PloS one, 15(9), e0239456. 10.1371/journal.pone.0239456

Perera, R. H., Abenojar, E., Nittyacharn, P., Wang, X., Ramamurthy, G., Peiris, P., Bederman, I., Basilion, J. P., & Exner, A. A. (2022). Intracellular vesicle entrapment of NB US contrast agents targeted to PSMA promotes prolonged enhancement and stability in vivo and in vitro. Nanotheranostics, 6(3), 270–285. 10.7150/ntno.64735

Przystupski, D., & Ussowicz, M. (2022). Landscape of Cellular Bioeffects Triggered by US-Induced Sonoporation. International journal of molecular sciences, 23(19), 11222. 10.3390/ijms231911222

Sarantis, P., Koustas, E., Papadimitropoulou, A., Papavassiliou, A. G., & Karamouzis, M. V. (2020). Pancreatic ductal adenocarcinoma: Treatment hurdles, tumor microenvironment and immunotherapy. World journal of gastrointestinal oncology, 12(2), 173–181. 10.4251/wjgo.v12.i2.173

Sharma, D., Giles, A., Hashim, A., Yip, J., Ji, Y., Do, N. N. A., Sebastiani, J., Tran, W. T., Farhat, G., Oelze, M., & Czarnota, G. J. (2019). Ultrasound microbubble potentiated enhancement of hyperthermia-effect in tumours. PloS one, 14(12), e0226475. 10.1371/journal.pone.0226475

Zhang, Y., Li, W., Niu, J., Fan, Z., Li, X., & Zhang, H. (2025). Reprogramming of glucose metabolism in pancreatic cancer: mechanisms, implications, and therapeutic perspectives. Frontiers in immunology, 16, 1586959. 10.3389/fimmu.2025.1586959

